# Potassium channel regulators are differentially expressed in hippocampi of Ts65Dn and Tc1 Down syndrome mouse models

**DOI:** 10.1101/467522

**Authors:** Shani Stern, Rinat Keren, Yongsung Kim, Elisha Moses

## Abstract

**Background:** Down syndrome remains the main genetic cause of intellectual disability, with an incidence rate of about 1 in 700 live births. The Ts65Dn mouse strain, with an extra murine chromosome that includes genes from chromosomes 10, 16 and 17 of the mouse and the Tc1 strain with an extra human chromosome 21, are currently accepted as informative and well-studied models for Down Syndrome. Using whole cell patch clamp we recently showed changes in several types of transmembrane currents in hippocampal neuronal cultures of Ts65Dn and Tc1 embryos. The associated genetic changes responsible for these changes in physiology were yet to be studied.

**Methods:** We used qPCR to measure RNA expression level of a few of the channel genes that we suspect are implicated in the previously reported changes of measured currents, and performed statistical analysis using Matlab procedures for the standard t-test and ANOVA and for calculating correlations between the RNA expression levels of several channel genes.

**Results:** We present differential gene expression levels measured using qPCR of the potassium channel regulators KCNE1 and KCNE2 in both Ts65Dn and Tc1 embryos and pups compared to controls. In Tc1, the human genes KCNJ6 and KCNJ15 are expressed in addition to a statistically insignificant increase of expression in the mouse genes KCNJ6 and KCNJ15. All channel genes that we have measured with large replication, have the same up-regulation or down-regulation in both mouse models, indicating that the transcription mechanism acts similarly in these two mouse models. The large dataset furthermore allows us to observe correlations between different channel genes. We find that, despite the significant changes in expression levels, channels that are known to interact have a high and significant correlation in expression both in controls and in the Down syndrome mouse model.

**Conclusions:** We suggest the differential expression of KCNE1 and KCNE2 as a possible cause for our previously reported changes in potassium currents. We report a KCNJ6 and KCNJ15 overexpression, which plays a role in the increased input conductance and the reduced cell excitability that we previously reported in the Tc1 mouse model. The large and significant positive (KCNQ2-KCNQ3, KCNE1-KCNE2, KCNQ3-KCNE1, KCNQ2-KCNE1, KCNQ2-KCNE2, KCNQ3-KCNE2) and negative correlations (KCNE1-KCNJ15, KCNE2-KCNJ15) that we find between channel genes indicate that these genes probably work in a cooperative or in a mutually exclusive manner.

## Introduction

Despite improved screening of embryos today, the rate of Down syndrome births has not decreased and is currently at a fixed rate of ^~^6000 babies annually in the USA alone [2]. The defined genetics of DS has allowed the development of good mice models, which have been shown to share behavioral phenotypes with human DS. In this study we used qPCR to measure six ion channel genes in the Tc1 and Ts65Dn mouse models. The Ts65Dn mouse model [3] is currently the most used and studied DS mouse model, with an extra translocation chromosome that includes genes from chromosome 16 and 17 of the mouse [4,5] and a total of about 50% of the DS genes in trisomy. Ts65Dn mice show developmental delays [6] and hyperactive behavior [3]. Ts65Dn mice were shown to have impaired LTP [7-10] as well as impaired learning in tasks such as the water maze [11] and other learning tasks [12]. The balance between excitation and inhibition was also shown to be altered [8]. Changes in synapse morphology were shown in the cortex and hippocampus [13], along with changes in neuronal densities in several brain areas such as the hippocampus, cortex and cerebellum [14,15]. A ^~^50% elevation in expression levels of the GIRK channel KCNJ6 were reported [16-18], and also of the IRK channel KCNJ15 [1,19]. A few studies have shown that blocking GABAA and GABAB inhibition improves deficits in cognition and synaptic plasticity [9,20-22].

The Tc1 mouse model [23] has an extra human chromosome 21 (^~^83% of the chromosome, [24]), and is mosaic with around 50% of the cells having the extra chromosome. At the behavioral level, the Tc1 model has decreased performance in cognitive and in locomotive tasks [23], impaired short term memory [25] and reduced exploratory behavior [26]. At the cellular level a reduction in cerebellar volume was reported [23], and altered protein profiles in the hippocampus [27].

We have recently used these 2 mouse models, Tc1 and Ts65Dn, to show a very clear phenotype of neuronal cultures that were derived from hippocampi of Tc1 and Ts65Dn embryos dissected at day 17 of the gestation period [1]. Network bursts had a lower amplitude and propagated slower in semi one dimensional cultures [28,29] for both mouse models, spike shape alterations and a smaller amplitude of fast AHP were reported for both mouse models. Hypoexcitability of the neurons were shown for both mouse models. Interestingly, a larger amplitude fast AHP was reported to be associated with hyperexcitability of hippocampal neurons in a study of bipolar disorder model neurons [30]. In both the Ts65Dn and Tc1 mouse models we found that the A-type and delayed rectifier potassium currents were reduced. Specific measurement in Ts65Dn cultures showed that HCN currents were increased by about 60%, while M-type currents were reduced by 90%. Up regulation of about 40% was observed for the expression of KCNJ15 in Ts65Dn, probably contributing to the neurons’ hypoexcitability. A computational model of the DS neuron yielded a very accurate reproduction of the experimental result.

Our measurements were generally performed under the assumption that the two potassium channel regulators KCNE1 and KCNE2, which reside in proximity to each other on chromosome 21, may have altered expression in DS. There are many reports that these channels interact with multiple potassium channels [31-39]. We therefore measured some of these currents that we hypothesized will change due to possible changes in KCNE1 and KCNE2 expression levels. Indeed, both our computational model and patch clamp recordings showed that these currents are altered. This leads us to examine KCNQ2 and KCNQ3, which are M-type channel genes for which both our computational model and patch clamp recordings indicate that the associated currents are altered [1]. Similarly, the KCNJ6 and KCNJ15 genes code for inward rectifying potassium channels whose presence in elevated quantities is essential to explain our previous observation of reduced excitability in DS neurons.

Here we measured these six genes that are associated with ion channel over a large number of pups and embryos, providing robust statistics. We find that KCNE2 is down-regulated in hippocampus of both mouse models. KCNE1 on the other hand is up-regulated in both mouse models. The KCNQ2 and KCNQ3 M-type channel genes are generally not affected, with a tendency to be up-regulated. The KCNJ6 and KCNJ15 mouse genes, which were shown by us and others to be up-regulated in the Ts65Dn hippocampus, have a similar expression in the Tc1 hippocampus with a tendency to be up-regulated, but in addition there is also expression of the human KCNJ6 and KCNJ15 channel genes. Observing the correlations between channel genes that were measured from the same samples reveal previously unknown correlations between genes that are known to interact and form complexes, giving further support to theoretical work on such correlations [40,41].

## Materials and Methods

Animal handling was done in accordance with the guidelines of the Institutional Animal Care and Use Committee (IACUC) of the Weizmann Institute of Science, and the appropriate Israeli law.

### Mice

Ts65Dn females were stock B6EiC3Sn a/A-Ts(1716)65Dn/J from Jackson catalog number 001924. The Ts65Dn strains are maintained by crossing Ts65Dn trisomic females with B6EiC3SnF1/J males. Ts65Dn litters comprise of Ts65Dn negative and Ts65Dn positive pups, and these are determined by genotyping. Original Tc1 females were received from the lab of Elizabeth Fisher at University College London, strain Tc1/(129S8xC57Bl/ 6)F1. These were mated with males of strain (129S8xC57Bl/6)F1, obtained by crossing C57Bl/6 males with females from stock 129S8/SvEvNimrJ, Jackson catalog 012809. The 129S8/SvEv strain was maintained by mating siblings. All embryos were used in each dissection performed, and all pups were taken in each litter that was used for the study. All animals were sacrificed in the lab with accordance to IACUC of the Weizmann Institute of Science.

Animal handling was done in accordance with the guidelines of the Institutional Animal Care and Use Committee (IACUC) of the Weizmann Institute of Science, and the appropriate Israeli law. The Weizmann Institute is accredited by AAALAC. The Weizmann Institutional Animal Care and Use Committee approved this study, conducted with mice hippocampi

### Genotyping

DNA was extracted using the Extract-N-Amp Tissue PCR kit from Sigma using ^~^5 mg of brain tissue or ^~^0.5 cm mouse tail. Genotyping of Ts65Dn was obtained using PCR with the (Reinholdt et al., 2011) protocol. For Tc1 mice we followed the protocol supplied courtesy of the Fisher lab, available in the Jackson homepage for genotyping of Tc1 (Jackson, 2010).

### RNA purification

For Ts65Dn mouse model, we analyzed mRNA of samples from embryos at day 17 of gestation (E17) and from newborn neonatal pups (P0). For Tc1 mouse model only P0 newborn neonatal pups were used. Hippocampi were harvested immediately after decapitation and dissection, frozen in liquid nitrogen and kept at -70C until the RNA extraction.

RNA was purified using the RNeasy Plus Micro kit (Qiagen, cat. 74034), and the protocol describing the purification is described in the RNeasy Plus Micro Handbook. After purification, the RNA was kept at –80^°^c until used. The amount of purified RNA was measured using the NanoDrop spectrophotometer. This step was performed completely without knowledge of embryo/pup genotype, and therefore is a blinding step.

### Reverse Transcription and amplification

Reverse transcription was performed using High-capacity cDNA Reverse transcription kit cat. 4368814 by ThermoFisher Scientific, using 2 μg of RNA per reaction. PCR was then performed using Extract-N-Amp PCR kit by Sigma-Aldrich, using the suggested protocol in the cDNA Reverse transcription kit protocol.

### QPCR

Quantitative RT-PCR was performed using the Applied Biosystems StepOne Plus system. Each RT-PCR reaction contained 5 μl of the Taqman gene expression master mix from Life Technologies (cat. no. 4369015), 0.5 μl of Taqman gene assays from Life Technologies, 2.5 μl RNAse-free water and 2 μl cDNA containing 100 ng cDNA. All samples were run in triplicates and the GAPDH gene (Mm99999915_g1, Life Technologies) was used as an endogenous control.

The gene assays that were used are the following. KCNJ15 mouse Mm02020346_s1, KCNJ15 human Hs01937913_s1, KCNJ6 mouse Mm01215650_m1, KCNJ6 human Hs01040524_m1, KCNE1 mouse Mm04207514_m1, KCNE2 mouse Mm00506492_m1, KCNQ2 mouse Mm00440080_m1, KCNQ3 mouse Mm00548884_m1, GAPDH Mm99999915_g1.

### Analysis

Analysis was done using the Matlab software package (2014b, The MathWorks Inc., Natick, MA, 2000). The expression level of the house keeping gene GAPDH was subtracted from the expression level of the gene that was examined in both DS and control Δ*C_T_* = *avg*(*C_T_*(*gene*)) − *avg*(*C_T_*(*GAPDH*)), *C_T_* being the PCR cycle at which the specified threshold level was crossed. RQ (Relative Quantification) of the control was taken as 1, while RQ of the DS population was taken as 2^Δ*C_T_*(*DS*)−Δ*C_T_*(*control*))^. The error for RQ was calculated for a 97% confidence interval using *RQ_min_* = 2^−2.14∗*ste*(*C_T_*(*gene*)−*C_T_*(*control*))^, and *RQ_min_* = 2^2.14∗*ste*(*C_T_*(*gene*)−*C_T_*(*control*))^, leading in general to asymmetric error bars. Statistical significance was tested using the student t-test between control Δ*C_T_* and DS Δ*C_T_*. The 2.14 in the exponent gives the 97% confidence interval.

## Results

### Down regulation of KCNE2 in both DS mouse models

KCNE2 is an important regulator of different voltage gated ion channels, which resides on chromosome 21. In a previous research [1], we hypothesized that changes to the expression of KCNE2 gene could cause the disrupted regulation of several types of potassium currents that we had observed, making it one of our target genes. Surprisingly instead of up regulation, we found a drastic down regulation. In the Tc1 mouse model the expression level in Tc1 pups and embryos was decreased by 66% in comparison with control pups and embryos (Fig. 1A, p=0.022, n=17 Tc1 pups and n=31 control pups). The expression level in Ts65Dn pups and embryos was decreased by 60% compared to control pups and embryos (Fig. 1B, p=0.049, n=40 Ts65Dn pups and n=44 control pups, using a power and sample size calculator with this change in the mean and with this large standard deviation showed that we need ^~^75 samples to achieve statistical significance, which is close to our sample size). Interestingly the diversity of the KCNE2 expression was much higher than the diversity of all the other channel genes we have measured in both mouse models and in their controls. The fold change varied in Tc1 mice from RQmin-RQmax of 0.097 to 1.183, and in control pups from 0.364 to 2.75. In the controls of the second mouse model it varied from 0.4839 to 2.0664, and in the Ts65Dn mice it varied from 0.2137 to 0.7622.

**Figure 1:**
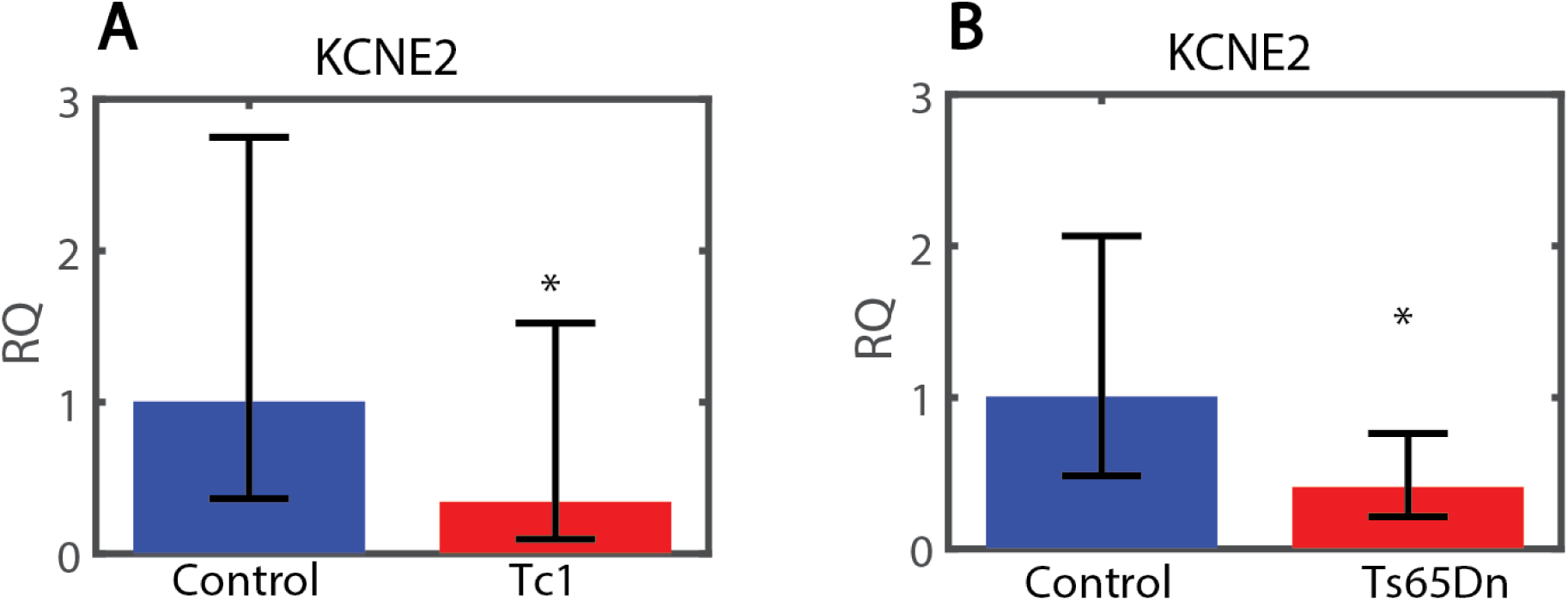
Expression levels of the KCNE2 gene in hippocampi of Tc1 and Ts65Dn DS mice models and their respective controls. **A.** KCNE2 level drops significantly in Tc1 mouse model compare to control littermates. **B.** KCNE2 level drops significantly in Ts65Dn mouse model compare to control littermates. Generally, the KCNE2 gene displays very large diversity in its expression profile for both control and DS model mice.

### Down regulation of KCNE2 in occurs in E17 Ts65Dn embryos, but not in P0s

When partitioning the expression data for KCNE2 to P0 vs. E17, we observe that the decrease in expression occurs in E17 Ts65Dn embryos, and to a much lesser extent, if at all, in P0s. In P0, a statistically insignificant decrease of 13% is observed in expression of KCNE2 in Ts65Dn pups vs. control littermates (Fig. 2A, p=0.82, n=23 Ts65Dn pups and n=24 control pups). In E17 embryos, a significant decrease of more than 80% is observed in KCNE2 expression in Ts65Dn embryos compared to their littermates (Fig. 2B, p=0.029, n=15 Ts65Dn embryos and n=17 control embryos).

**Figure 2:**
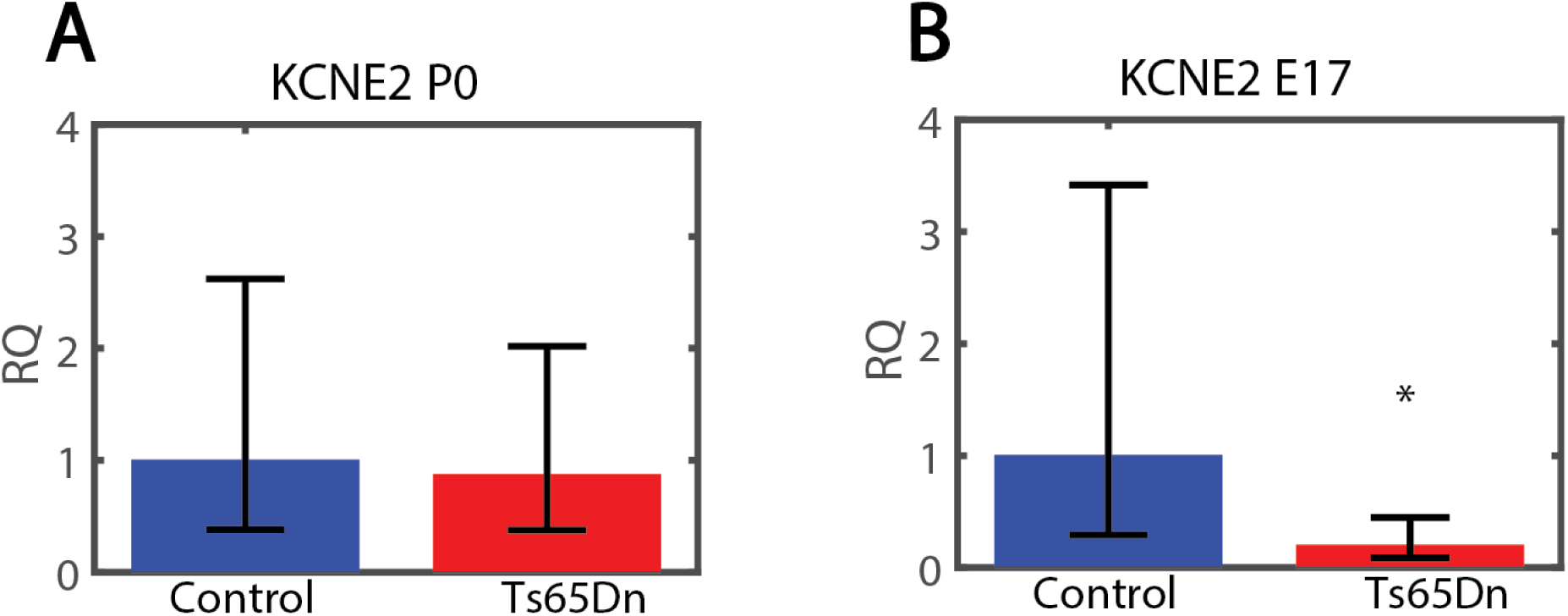
Expression levels of KCNE2 gene in hippocampi of P0 pups and of E17 embryos of Ts65Dn mice vs. control littermates. **A.** KCNE2 expression in Ts65Dn P0 pups shows is similar to that in controls. **B.** KCNE2 is down regulated in Ts65Dn embryos compared to their control littermates.

### Up regulation of KCNE1 in both DS mouse models

KCNE1 is also an important regulator of different voltage gated ion channels, which resides on chromosome 21, less than 80000 base pairs away from KCNE2. In our previous paper [1], we hypothesized that changes to the expression of KCNE1 would also be involved in dysregulation of potassium currents. Indeed we report some up regulation of this channel in both mouse models, although this up regulation was not statistically significant. In the Tc1 mouse model the expression level in pups was ^~^50% higher than in control pups (Fig. 3A, p=0.097, n=16 Tc1 mice, n= 30 control mice). In the Ts65Dn mouse model the expression level in Ts65Dn pups and embryos was similarly about 50% higher than in control pups and embryos (Fig. 3B, p=0.14, n=41 Ts65Dn mice, and n=39 respective controls).

**Figure 3:**
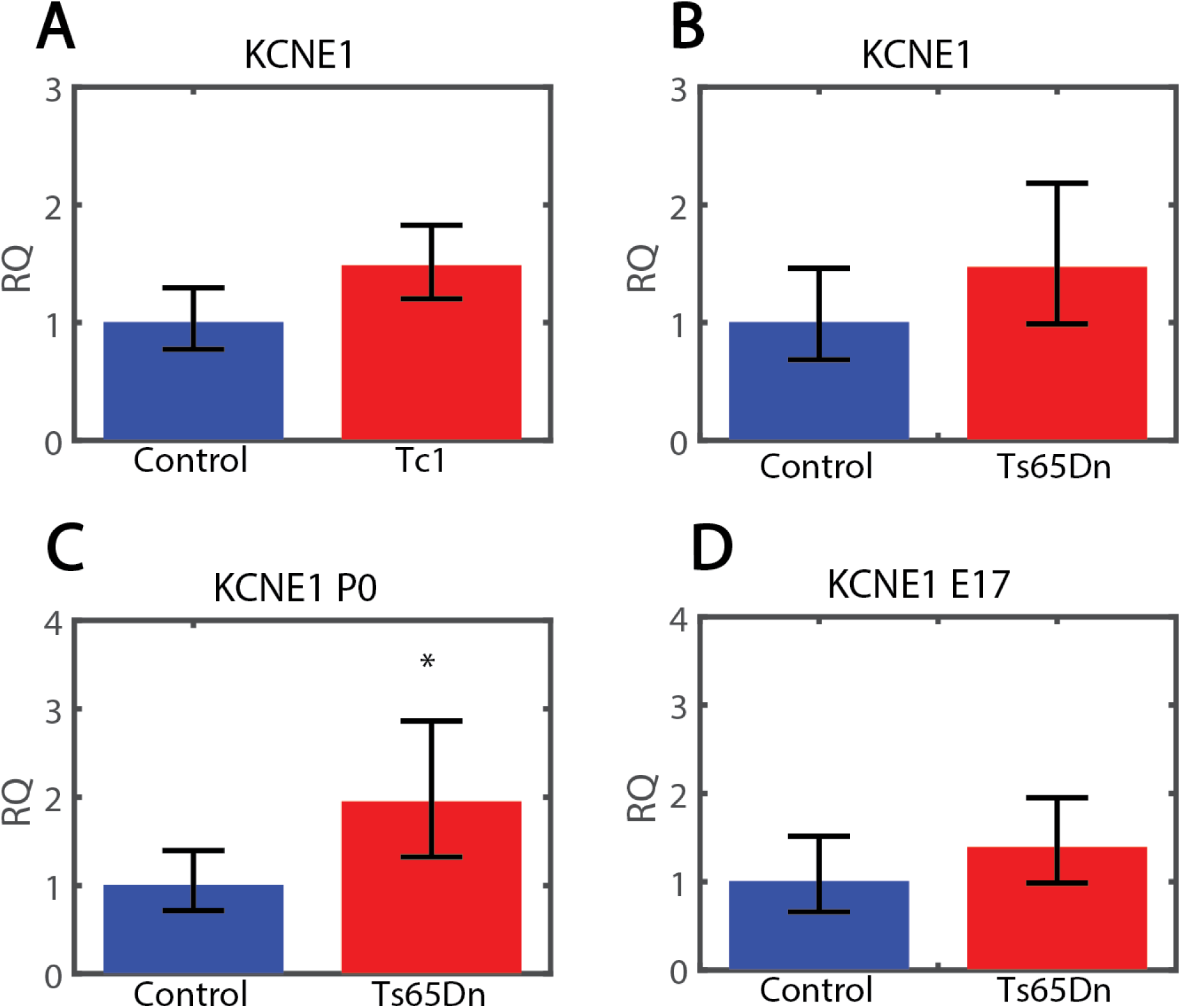
Expression levels of the KCNE1 gene in hippocampi of Tc1 and Ts65Dn DS mice models and their respective controls. **A.** KCNE1 level is elevated in Tc1 mouse model compare to control littermates. **B.** KCNE1 level is elevated in Ts65Dn mouse model compare to control littermates. **C.** KCNE1 level is elevated in Ts65Dn P0 pups mouse model compared to control littermates. **D.** KCNE1 is not significantly elevated in Ts65Dn E17 embryos compared to their control littermates.

### Up regulation of KCNE1 is significant in P0 pups, but not in E17 embryos

When partitioning the data to P0 vs. E17, we observe that the increase in the KCNE1 expression is significant only in Ts65Dn P0 pups, and occurs to a much lesser extent in Ts65Dn E17 embryos). An increase of 95% in expression of KCNE1 gene is observed in Ts65Dn P0 pups compared to their littermate controls (Fig. 3C, p=0.016, n=6 Ts65Dn P0s, n=8 control P0s). In E17 embryos, an increase of 39% (not significant) is observed in the expression level of KCNE1 (Fig.3D, p=0.19, n=30 Ts65Dn E17 embryos, n=26 control E17 embryos).

### KCNQ2 shows a tendency to be over-expressed in DS mice, but not significantly

In a previous paper [1], we showed that M-type currents are almost completely abolished in Ts65Dn hippocampal cultures. We therefore measured expression levels of two of the genes associated with M-type currents, KCNQ2 and KCNQ3. KCNQ2 does not show any changes in expression in either Tc1 or Ts65Dn mouse models, with a tendency to be upregulated. In Tc1 P0 pups there is a statistically insignificant increase of 14% in expression of KCNQ2 compared to P0 control pups (Fig. 4A, p=0.39, n=14 Tc1 pups, n=28 control pups). In Ts65Dn E17 embryos and p0 pups there is practically no change (increase of 8%) in KCNQ2 expression compared to control E17 embryos and pups (Fig. 4B, p=0.37, n=23 Ts65Dn embryos and pups, n=26 control embryos and pups). There is no significant change in expression between E17 embryos and P0 pups in the KCNQ2 gene (Supplementary Fig. 1)

**Figure 4:**
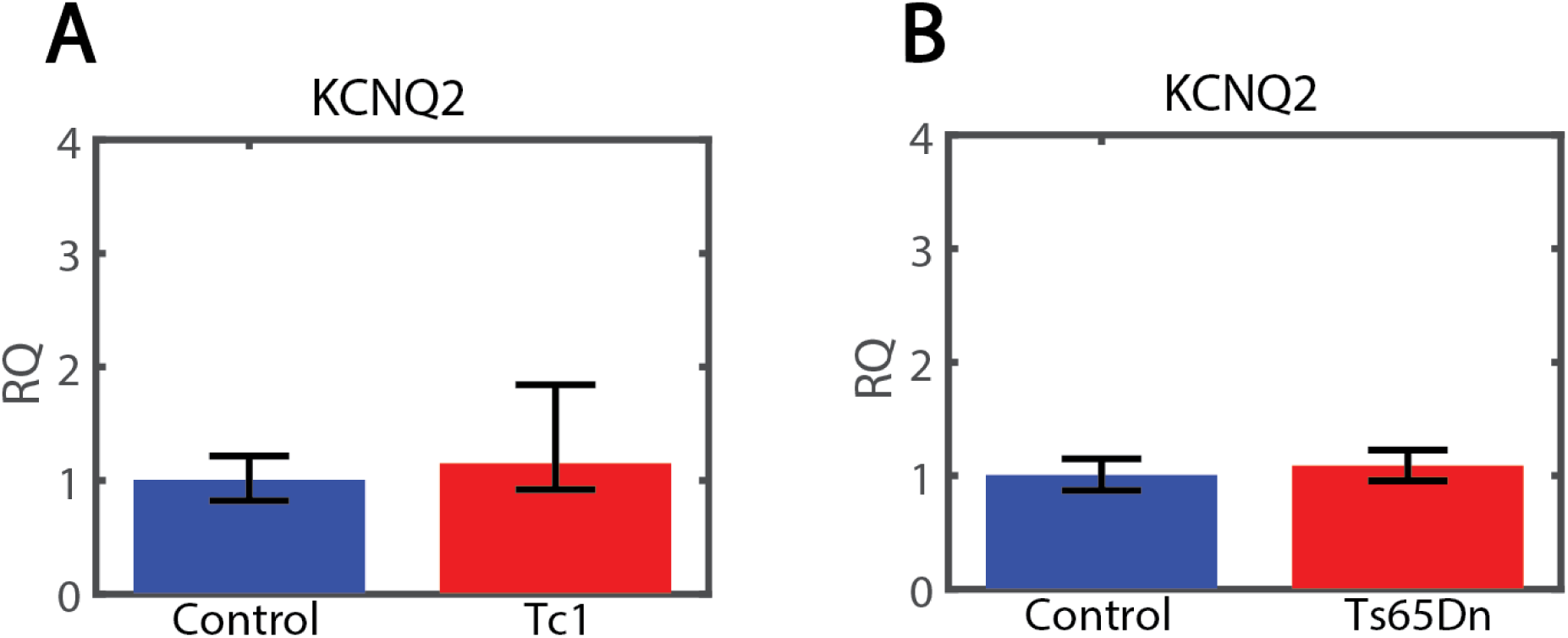
Expression levels of KCNQ2 gene in hippocampi of Ts65Dn mice and Tc1 mice compared to control mice. **A.** KCNQ2 level is slightly elevated in Tc1 pups compared to their control littermates (N.S.). **B.** KCNQ2 level is slightly elevated in Ts65Dn pups compared to their control littermates (N.S.).

### KCNQ3 shows a tendency to be over-expressed in DS mice, but significance is achieved only when pulling together all DS mice together

KCNQ3 does not show a significant change in expression in both Tc1 and Ts65Dn mouse models, but the overall expression level becomes significantly different in DS when pooling together data from both DS mouse models. In Tc1 P0 pups there is an increase of 30% in KCNQ3 gene expression compared to control P0 pups (Fig. 5A, p=0.13, n=14 Tc1 P0 pups, n=28 control P0 pups). In Ts65Dn P0 pups there is an increase of 39% compared to control P0 pups (Fig. 5B, p=0.24, n=21 Ts65Dn p0 pups, n=22 control p0 pups). When combining the two DS mouse models together, there is an increase of 51% in DS P0 pups compared to control E17 p0 pups (Fig. 5C, p=0.026, n=35 DS P0 pups, n=50 P0 control pups).

**Figure 5:**
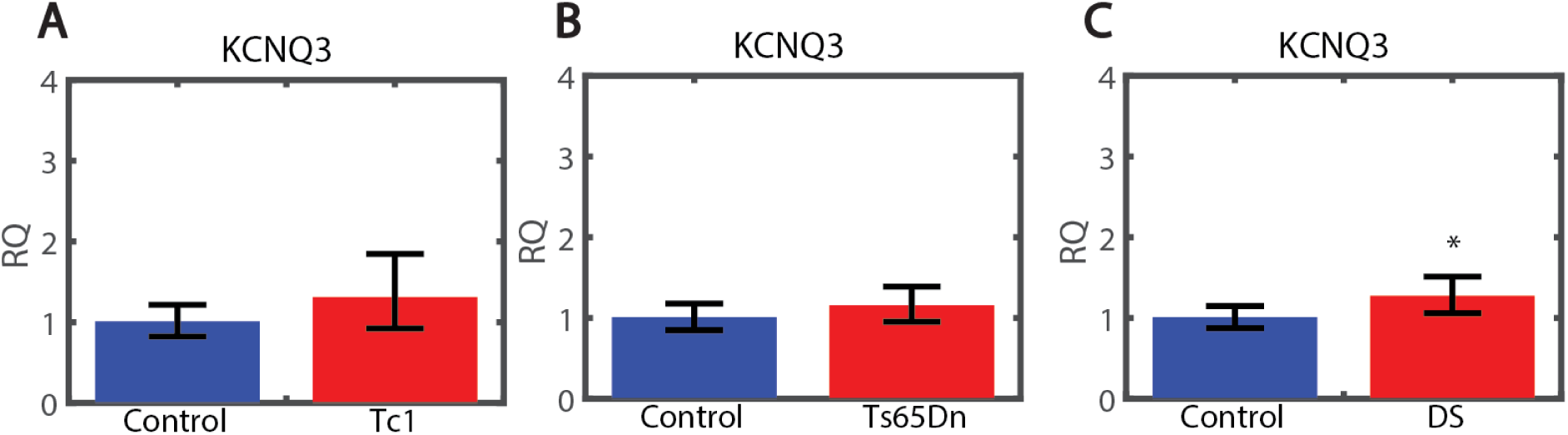
Expression levels of KCNQ3 gene in hippocampi of Ts65Dn mice and Tc1 mice compared to control mice. **A.** KCNQ3 level is elevated (N.S.) in Tc1 pups compared to their control littermates. **B.** KCNQ3 level is slightly elevated in Ts65Dn pups compared to their control littermates (N.S.). **C.** When combining all DS data together, there is a significant increase in KCNQ3 expression of DS model mice compared to their control littermates.

### Upregulation of channel genes in P0 pups compared to E17 embryos in control and Ts65Dn mice

In view of the possibly different expression of the channel genes in P0 pups versus E17 embryos, we summarize in Figure 6 data from the control and the Ts65Dn model mice for all four channel genes. Since we expect significant brain development to occur between E17 and P0, we hypothesize that some ion channel genes may have different expression levels in P0 brains. SHANI - SAY WHY YOU DON’T SHOW Tc1.

**Figure 6:**
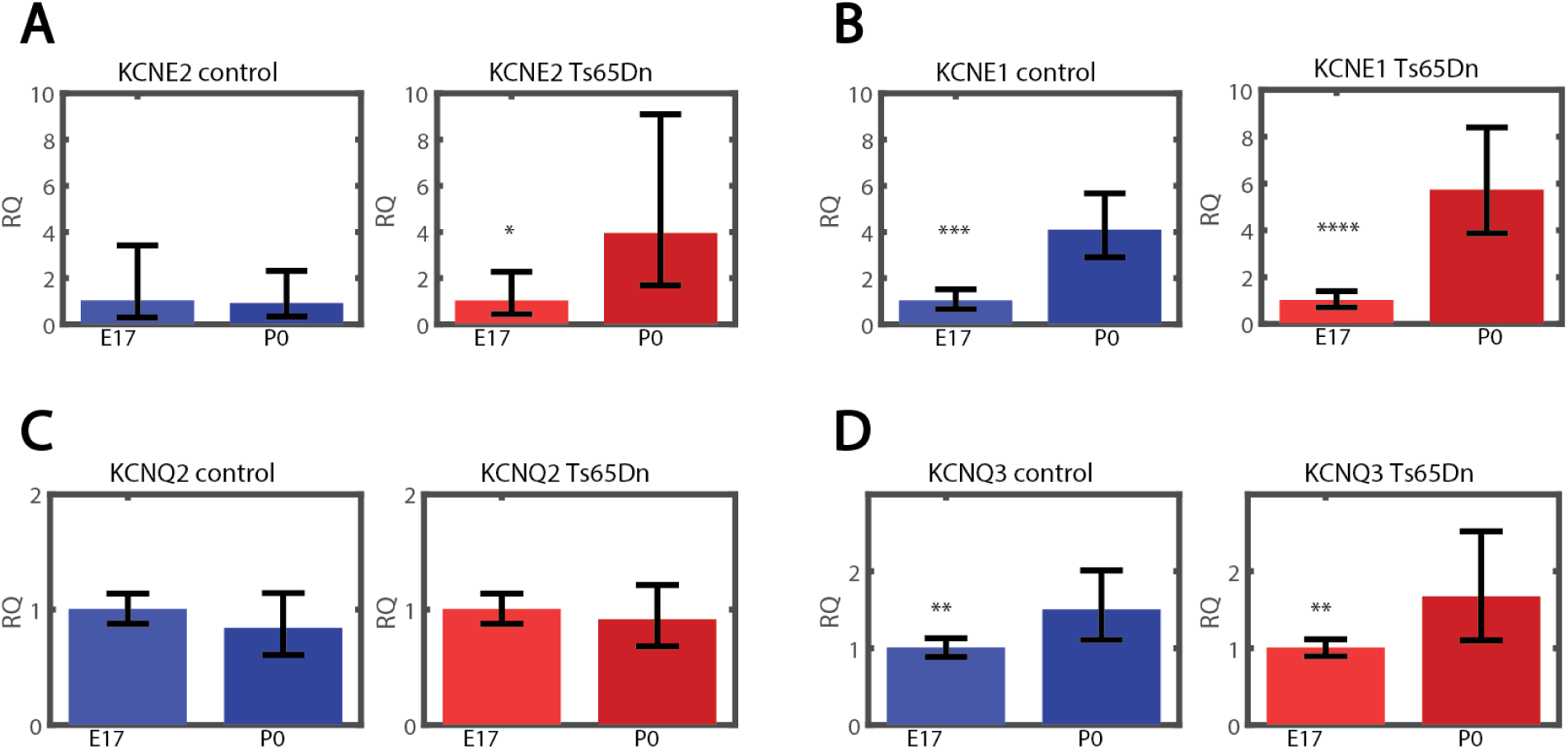
Expression levels of KCNE2, KCNE1, KCNQ2 and KCNQ3 genes in hippocampi of P0 pups vs. of E17 embryos in Ts65Dn mice and in control littermates. **A. Left:** KCNE2 expression in control P0 pups is similar to E17 embryos. **Right:** KCNE2 has a larger expression in Ts65Dn P0s compared to E17 embryos. **B. Left:** KCNE1 level is highly elevated in control P0 pups mouse model compared to E17 embryos. **Right:** KCNE1 level is highly elevated in Ts65Dn P0 pups compared to E17 embryos. **C. Left:** In control mice KCNQ2 expression in P0 pups is similar to E17 embryos. **Right:** In Ts65Dn mice KCNQ2 expression is also similar between P0 pups and E17 embryos. **D. Left:** In control mice KCNQ3 expression in P0 pups increases compared to E17 embryos. **Right:** In Ts65Dn mice KCNQ3 expression in P0 pups also increases compared to E17 embryos.

For KCNE2 gene, expression is slightly (^~^12%) decreased in control P0 pups compared to control E17 embryos (p=0.86, n=24 control P0 pups, n=17 control E17 embryos, Fig. 6A left panel). In contrast, expression is increased almost three-fold in Ts65Dn P0 pups compared to E17 embryos (p=0.023, n=23 Ts65Dn P0 pups, n=15 Ts65Dn E17 embryos, Fig. 6B right panel).

The KCNE1 gene is also differentially expressed in P0 pups compared to E17 embryos, supporting further the hypothesis of a developmental stage of the CNS and consequent differential expression of ion channels. However, in this gene, we find a large difference in both control and Ts65Dn mice. In control P0 pups, KCNE1 is three-fold more expressed than in control E17 embryos (306% increase, p=0.0005, n=8 control P0 pups, n=26 control E17 embryos, Fig. 6B left panel). In Ts65Dn P0 pups, KCNE1 is almost fivefold more expressed than in E17 embryos (471% increase, p=0.00004, n=6 Ts65Dn P0 pups, n=30 Ts65Dn E17 embryos, Fig. 6B, right panel). It should be noted that both the KCNE1 and KCNE2 genes are in trisomy in Ts65Dn, possibly accounting for the more pronounced increase in these genes expression levels between P0 and E17 is observed.

KCNQ2 is not differentially expressed between P0 pups and E17 embryos in control and Ts65Dn mice. In control mice KCNQ2 expression in P0 pups is 17% reduced compared to E17 embryos (p=0.2, n=9 P0 pups, n=17 E17 embryos, Fig. 6C, left panel). In Ts65Dn mice KCNQ2 expression in P0 pups is 8% reduced (p=0.47, n=7 P0 pups, n=16 E17 embryos, Fig. 6C, right panel).

Finally, the KCNQ3 gene is also differentially expressed in P0 pups compared to E17 embryos in both control and Ts65Dn mice. In control mice KCNQ3 expression in P0 pups is increased by almost 50% compared to E17 embryos (p=0.007, n=7 P0 pups, n=13 E17 embryos, Fig. 6D, left panel). In Ts65Dn mice KCNQ3 expression in P0 pups is increased by 66% (p=0.0035, n=7 P0 pups, n=14 E17 embryos, Fig. 6D, right panel).

We conclude that between E17 embryos and P0 pups, 3 out of these 4 ion channel genes exhibit a clear increase in expression. The most pronounced effect is in the KCNE1 gene, even though its expression in the hippocampi is generally low.

### Mouse KCNJ6 shows slightly elevated expression in Tc1 mice, and in addition there is expression of human KCNJ6 gene in Tc1 mice

It has previously been reported by others that the KCNJ6 gene is overexpressed in Ts65Dn E17 embryos [16]. We assessed the over-all expression level of KCNJ6 in the Tc1 DS mouse model. The expression level of KCNJ6 in Tc1 P0 is increased by 14% (N.S.) compared to their respective control littermates (Fig. 7A, p=0.5 n=13 Tc1 P0 pups, n=22 control P0 pups). The Tc1 pups however also express human KCNJ6 gene, while control P0 do not express it. There is a ^~^400 fold expression in human KCNJ6 expression in Tc1 pups compared to control littermates (Fig. 7B, p=3e-25, n=13 Tc1 P0 pups, n=24 control P0 pups). The expression of human KCNJ6 is negligible and is due to imperfect specificity of the primer (and therefore the 400 fold change in expression is somewhat arbitrary. Our results simply indicate that the Tc1 DS mouse model expressed both the mouse and human gene, while the control mouse expresses only the mouse gene).

**Figure 7:**
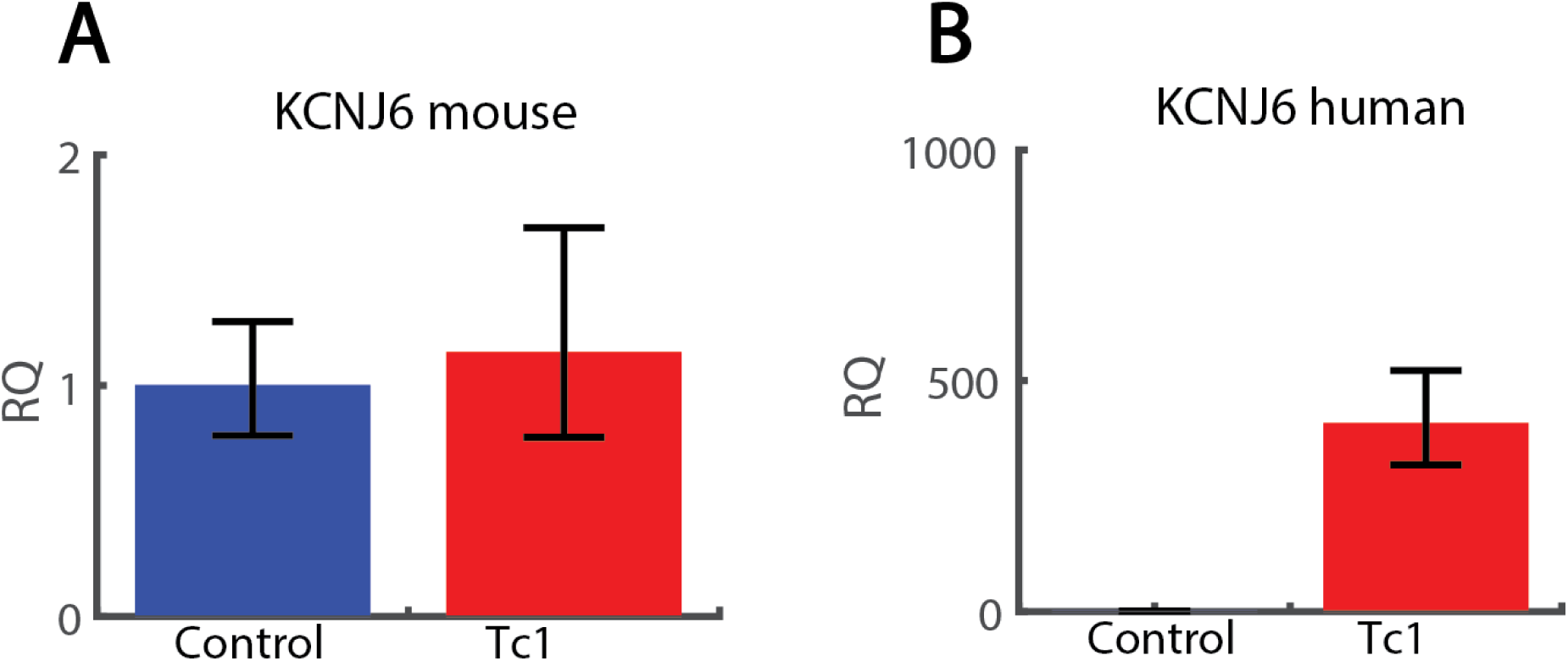
Expression levels of mouse and human KCNJ6 gene in hippocampi of Tc1 mice compared to control mice. **A.** KCNJ6 level is elevated (N.S.) in Tc1 pups compared to their control littermates. **B.** In addition to mouse KCNJ6, there is expression in Tc1 mice of human KCNJ6 gene from the human chromosome 21. In control mice the expression of human KCNJ6 is negligible and is due to imperfect specificity of the primer.

### Mouse KCNJ15 gene shows slightly elevated expression in Tc1 mice, and in addition there is expression of human KCNJ15 gene in Tc1 mice

In our previous report [1] we showed that KCNJ15 gene is overexpressed in Ts65Dn E17 embryos. We now assess the over-all expression level of KCNJ15 in the Tc1 DS mouse model. The expression level of KCNJ15 in Tc1 P0 is increased by a statistically insignificant 27% compared to their respective littermates (Fig. 8A, p=0.28 n=13 Tc1 P0 pups, n=22 control P0 pups). The Tc1 pups however also express human KCNJ15 gene, while control P0 do not express it. There is a ^~^10^4^-fold expression in human KCNJ15 expression in Tc1 pups compared to control littermates (Fig. 8B, p=2e-13, n=13 Tc1 P0 pups, n=24 control P0 pups). The expression of human KCNJ15 is negligible and is due to imperfect specificity of the primer (again the 10^4^ fold change in expression is arbitrary and our results indicate that the Tc1 DS mouse model expresses both the mouse and human gene, while the control mouse expresses only the mouse gene).

**Figure 8:**
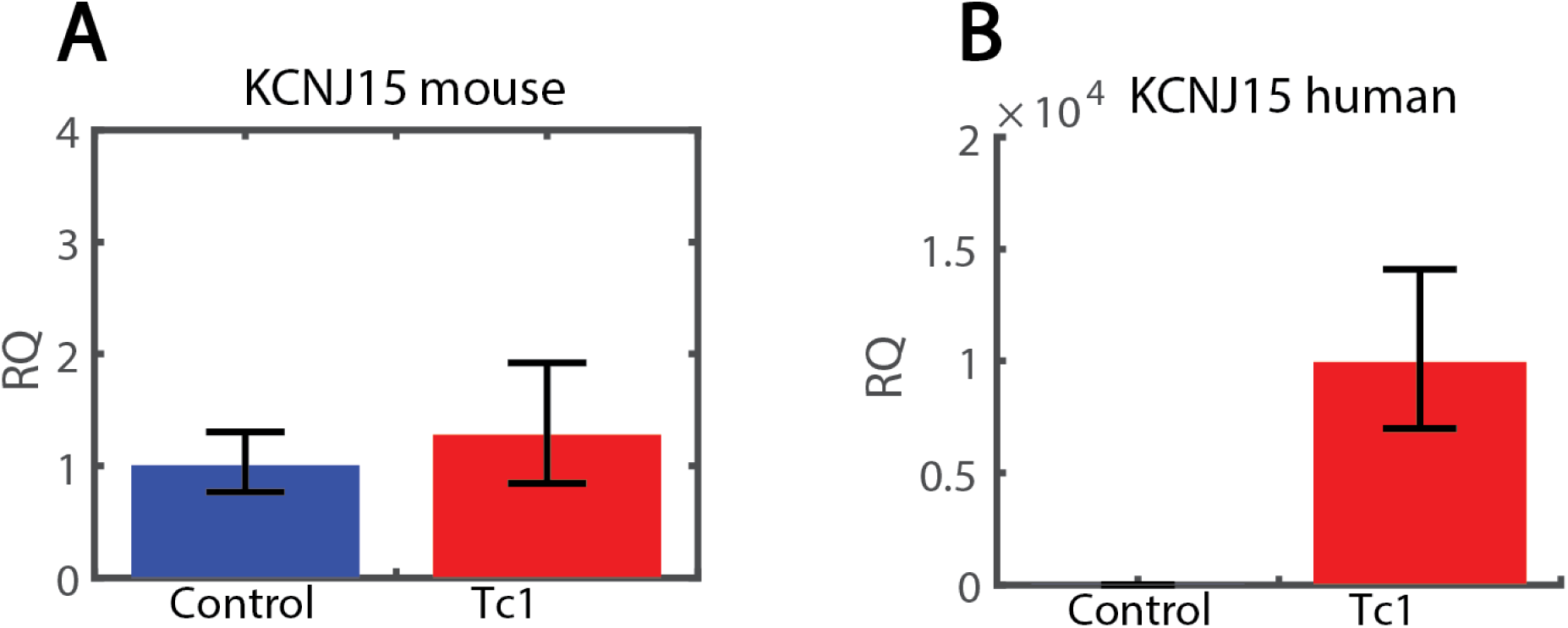
Expression levels of mouse and human KCNJ15 gene in hippocampi of Tc1 mice compared to control mice. **A.** KCNJ15 level is elevated (N.S.) in Tc1 pups compared to their control littermates. **B.** In addition to mouse KCNJ15, there is expression of human KCNJ15 gene from the human chromosome 21. The expression of human KCNJ6 in control mice is negligible and is due to imperfect specificity.

### Correlations of channel genes

We analyzed the correlations between expression levels for all pairs of genes, except those that link the KCNQ family with the KCNJ one, for which we had low statistics. We found no difference in these correlation between Ts65Dn, Tc1 and controls, and we therefore aggregate the data for all three strains. Remarkable, despite the differences in expression that are observed between the strains, the correlations between the genes are significant and alike across the strains.

Interestingly, we see strong correlations that were not reported previously. KCNQ2 and KCNQ3 are two genes that encode for M-type channels, and are known to work in cooperation to form M-type channels [42,43]. We found that, although they reside on different chromosomes, the expression level of these genes is highly correlated, R=0.9109, p<1e-10 (Fig. 9A). KCNE1 and KCNE2 reside on the same chromosome and are separated by less than 80000 base pairs. In addition, both KCNE1 and KCNE2 are known to interact with several types of potassium channels [31-39].The correlation that we measured between these two channel genes is R=0.62, p=1.9e-04 (Fig. 9B). The correlation that we measured between KCNE1 and KCNQ3 was R=0.7412, p=3.1e-06 (Fig. 9C). The correlation between expression of KCNE1 and KCNQ2 was R=0.7324, p=4.3e-06 (Fig. 9D). The KCNE2-KCNQ2 expression correlation is R=0.6514, p=1.2e-5 (Fig. 9E). The KCNE2-KCNQ3 expression correlation is R=0.7087, p=1.2e-6 (Fig. 9F). Such correlations are particularly surprising given the high diversity exhibited in the expression of KCNE2.

**Figure 9:**
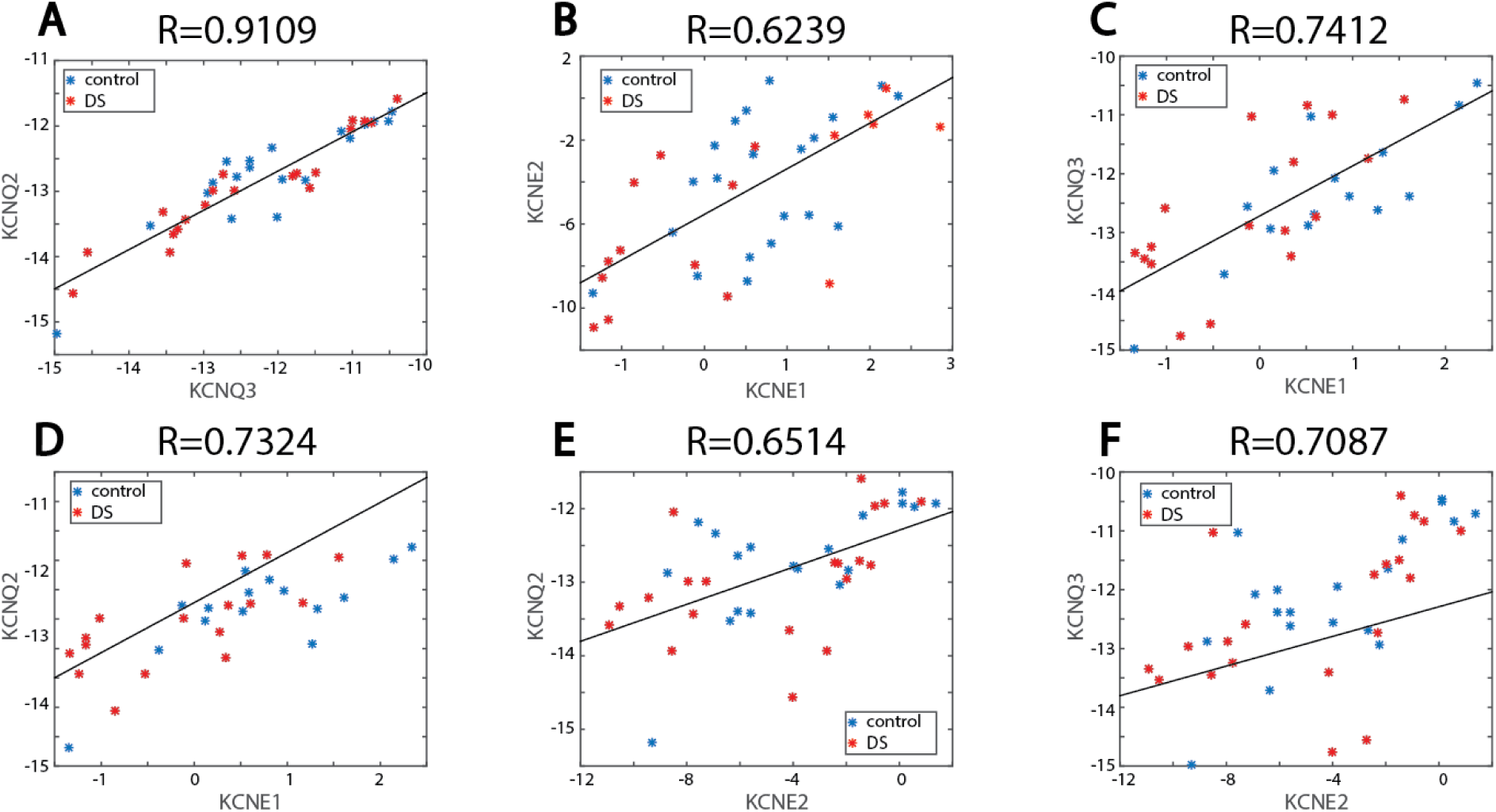
Correlation between gene expression levels of the different channel genes that we measured. The x and y axis are the Δ*C_T_* of each of the genes presented. (**A.** Strong positive and significant correlation between KCNQ2 and KCNQ3. **B.** Significant positive correlations between KCNE1 and KCNE2. **C.** Significant positive correlations between KCNE1 and KCNQ3. **D.** Significant positive correlations between KCNE1 and KCNQ2. **E.** Significant positive correlations between KCNE2 and KCNQ2. **F.** Significant positive correlations between KCNE2 and KCNQ3

There were no obvious correlations between other pairs of genes, with some surprising negative correlations. The KCNJ6-KCNE1 expression correlation was R=0.2979, p=0.13 (Fig. 10A). The KCNJ6-KCNE2 expression correlation was R=-0.1665, p=0.37 (Fig. 10B). The KCNJ6-KCNJ15 expression correlation was 0.2513, p=0.16 (Fig. 10C). Surprisingly KCNE1 and KCNE2 are negatively correlated with KCNJ15. KCNE1-KCNJ15 expression correlation was R=-0.45, p=0.01 (Fig. 10D). The KCNE2-KCNJ15 expression correlation was R=-0.4040, p=0.025 (Fig. 10E).

**Figure 10:**
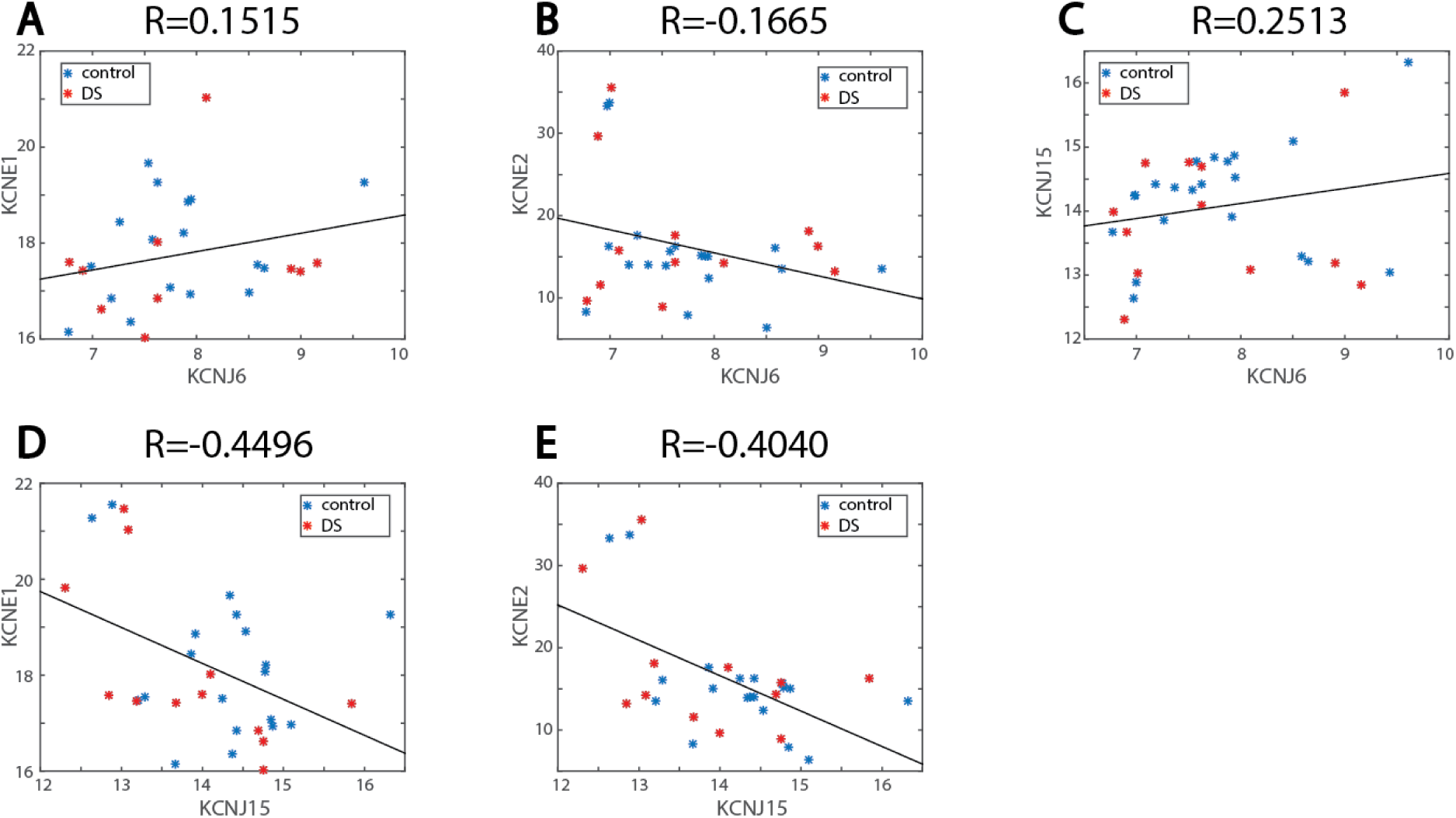
Absence of statistically significant correlation between expression levels of some different channel genes that we measured. **A.** No significant correlation is observed between KCNJ6 and KCNE1. **B.** No significant correlation is observed between KCNJ6 and KCNE2. **C.** No significant correlation is observed between KCNJ6 and KCNJ15. **D.** A negative correlation is measured between KCNJ15 and KCNE1. **E.** A negative correlation is measured between KCNJ15 and KCNE2.

### High expression noise in KCNE2

One of the interesting phenomena that we see from the data is a very noisy expression of KCNE2, and to a much lesser extent also of KCNE1. Figure 11 presents the standard variance (the expectation of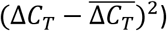
of the six channel genes. The noisy expression of KCNE2 was already reported in cortices of mice embryos in [44]. Here we report it for hippocampi. Using the F-test for equality of variances
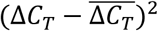
we tested for different variances with multiple hypotheses correction gives a p=1.2e-33 as the probability that the variances are similar between the expression levels of these channel genes. Note that the results are given in number of cycles of the qPCR. This means that the actual difference in expression level is 2 to the power of the standard deviation, i.e. the changes are exponential in the variance.

**Figure 11:**
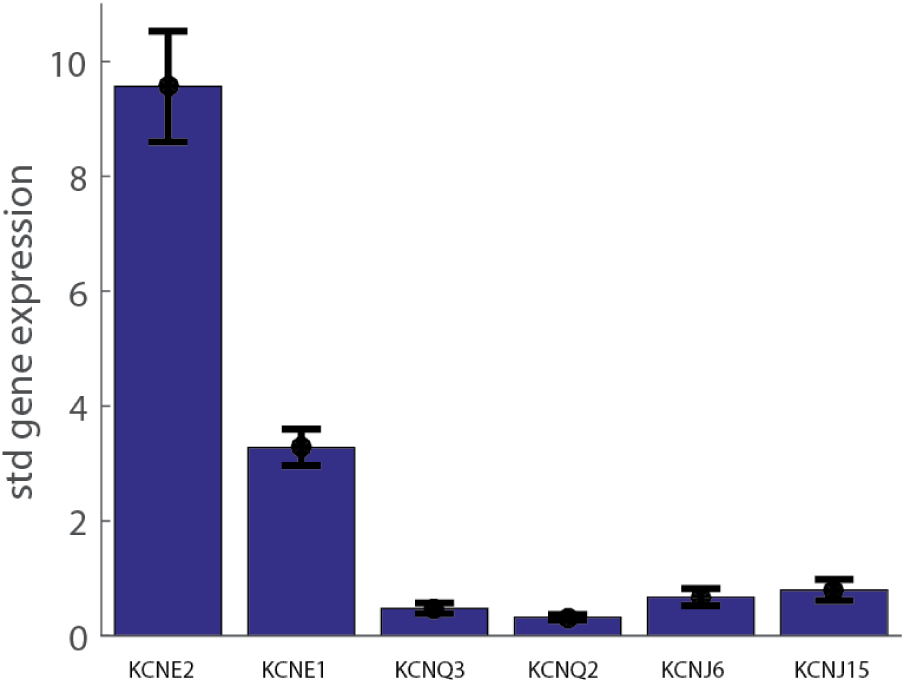
Variance in ***C_T_*** of the six channel genes. A very large variance is observed for KCNE2, and to a lesser extent of KCNE1, indicating that these channels have a noisy RNA expression level. The variance is calculated as
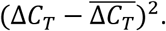

## Conclusions

We previously showed [1] that conductance of several potassium channels is altered in neuronal cultures derived from hippocampi of Ts65Dn and Tc1 embryos. Here we investigated how changes in expression levels of six associated ion channel genes may be involved in these changes. The genes we focus on that reside on chromosome 21 are the KCNJ6 and KCNJ15 ion channel genes and KCNE1 and KCNE2, which are potassium channel regulators. In addition we were specifically interested in the KCNQ2 and KCNQ3 genes of channels that comprise the M-type currents, since we have shown that the M-type currents are 90% reduced in hippocampi of Ts65Dn embryos.

While there are a few theoretical results [40,41] which predict that channel genes which act together will be co-expressed, there are only a few experimental work supporting them [45-48]. Here we show correlations that were not yet reported of expression levels for several ion channel genes that are known to co-assemble and work cooperatively.

We have shown that the potassium channel regulators KCNE1 and KCNE2 indeed have altered regulation in Ts65Dn and Tc1 hippocampi in comparison to control littermates. KCNE1 and KCNE2 share 34% sequence identity and are thought to originate from gene duplication, and to be related through divergent evolution [49]. Both were shown to be voltage-gated potassium channel accessory subunits (beta-subunits) and have the ability to regulate multiple different ion channels [31-39] by co-assembling with alpha subunits. Interestingly, although these channels are neighboring genes, the KCNE2 gene is down-regulated both in Ts65Dn and Tc1 embryos and pups hippocampi, while KCNE1 is up-regulated both in Ts65Dn and Tc1 hippocampi. Furthermore, the expression pattern of both KCNE1 and KCNE2 is noisy (Fig. 11), albeit much noisier in KCNE2. This enhanced variability was previously observed in cerebellum, cortex and midbrain [44]. KCNE1 is strongly up-regulated between E17 and P0 pups, suggesting that this is an important developmental protein, shaping neuronal maturation and function. With KCNE2 this is less clear, due to its inherently large noise, but it also appears to be strongly up-regulated between E17 and P0. The effect in both regulators is stronger in the Ts65Dn embryos and pups than in the controls, and we hypothesize that this is due to the extra copy of the chromosome in DS. It should be noted, that although KCNE1 and KCNE2 reside on chromosome 21, the Tc1 mouse model does not have 3 copies of these genes [24].

We have seen that KCNQ2 and KCNQ3 in our data tend to be up-regulated, with a statistically significant result of about 50% increase in expression obtained only when pooling all the data together in the KCNQ3 gene. In a previous publication [1], the M-type currents were almost completely abolished in hippocampal neurons of Ts65Dn embryos. There are two possible explanations for the fact that we do not see a concurrent reduction in gene expression. The first is that the strong reduction that we observe in KCNE2 gene expression of both DS model mice (Fig. 1) may play a strong role in the reduction of the M-type currents [36,42]. The second possibility is that the electrophysiological measurements in our previous study were performed after 13-14 days of culturing the neurons, while the expression level is measured immediately after dissection. It may be that there is a faster degeneration of M-type channels in neurons derived from DS model mice compared to the control mice, which may relate to a general phenomenon of age related degeneration in DS that is known from Ts65Dn and human DS [50].

Mouse KCNJ15 and KCNJ6 genes were previously shown by us [1] and others [16] to be up-regulated in the hippocampus of Ts65Dn embryos. Here we show that there is practically the same expression of the mouse genes in the Tc1 P0 pups compared to their control littermates, but there is also expression of the human KCNJ6 and KCNJ15, which is of course completely missing from the control pups. This overabundance of inward rectifier channels may be the explanation for the reduced excitability that we have reported previously in neurons derived from Tc1 mouse embryos. This reduction manifested in smaller amplitude and shorter duration network bursts, and in fewer evoked action potentials in response to current injection.

It is interesting to note that in our previous report [1] all functional measurements that were performed for both mouse models showed a similar phenotype. This clearly indicates that despite the very different genetic modeling of DS, these two mouse models share many phenotypic features. Similarly, in the present study, all channels that were measured show the same tendency for up or down regulation in both the DS mouse models. It thus seems that these two mouse models have many properties in common. In fact we have also recently reported on the gradual abolishment of transmission of trisomy as the female ages, with very similar statistics in both mouse models [51].

In general, the 50% over-expression expected in trisomy of DS genes is difficult to detect with qPCR. We had to collect rather large statistics (a total of 137 pups and embryos for the noisier channel genes KCNE1 and KCNE2). These large statistics further allowed us to observe correlations between levels of expression of channel genes. There are some theoretical work [40,41] predicting that ion channels that interact with each other will be correlated in the expression level, but relatively few experimental results have been reported [45-48]. A very striking correlation between KCNQ2 and KCNQ3 expression of 0.91 was found with p<1e-10. These channels reside on different chromosomes, but are known to form channels together [43,52]. It is furthermore reported that the promoters of these genes share common elements [52]. Other interesting correlations that we found are KCNE2-KCNE1 (R=0.62), KCNQ3-KCNE1 (0.74), KCNQ2-KCNE1 (0.73), KCNQ3-KCNE2 (0.71) and KCNQ2-KCNE2 (0.65). These correlations are impressively high relative to other measurements of correlations between channel expression, and are all obtained for channels that are known to interact. As expected, there is generally no correlation between the KCNE1 and KCNE2 and the inward rectifiers KCNJ6 and KCNJ15. A negative correlation was found between KCNE1 and KCNJ15 (-0.45), and KCNE2 and KCNJ15 (-0.4), but the significance is not as strong as with the other correlations (p=0.01 and p=0.025 respectively). Looking at the graph (Fig. 10D and Fig. 10E) it is not immediately clear if indeed there is a negative correlation. But if a true negative correlation persists in future measurements, it predicts the tantalizing possibility that these genes may have antagonist functions and will not co-express.

To summarize, we have presented expression profiles of six ion channel genes in hippocampi of Ts65Dn, Tc1 and control embryos and pups. As suggested by our previous publication [1], KCNE1 and KCNE2 play a role in shaping changes in neuronal function in neurons from hippocampi of the two major yet completely different DS mouse models. We further show that the inward rectifying channel genes KCNJ6 and KCNJ15 have an overall up-regulation in the Tc1 mouse model, in line with previous reports on the Ts65Dn mouse model. The overall regulation pattern we observe is very similar between the two mice models for all six ion channel genes, lending confidence to our results and to the models themselves. We further showed the existence of correlations in expression of ion channel genes that interact that were not previously reported.

## Acknowledgments

The authors would like to thank Menahem Segal for very useful discussions. The authors thank Elizabeth Fisher for supplying the Tc1 mice and for many helpful suggestions and remarks. This work was partly supported by the Minerva Foundation (Munich, Germany) and by the Israel Science Foundation grant 1415/12 and the Clore Center for Biological Physics.

